# Spontaneous biases enhance generalisation in the neonate brain

**DOI:** 10.1101/2023.07.11.548572

**Authors:** Shuge Wang, Vera Vasas, Laura Freeland, Daniel Osorio, Elisabetta Versace

## Abstract

The ability to use sparse evidence to produce adaptive responses in new contexts and to new stimuli (inductive generalisation) is central to biological and artificial intelligence. Young and inexperienced animals require very little evidence to generalise, raising the question of whether the neonate brain is evolutionarily prepared (predisposed) for generalisation. To understand the principles of spontaneous generalisation, we exposed neonate chicks to an artificial social partner of a specific colour, and measured generalisation by comparing responses to novel and familiar stimuli along either the red-yellow or the blue-green colour continuum. Generalisation responses were inconsistent with an unbiased model, showing biases such as asymmetrical generalisation gradients, faster learning for particular colours (red and blue over yellow and green), preferences for unfamiliar stimuli and different time courses in learning. Moreover, the chicks’ generalisation behaviour was consistent with a Bayesian theoretical model that explicitly incorporates predispositions as initial preferences and treats the learning process as an update of spontaneous preferences. These results show that neonate animals are evolutionarily prepared for generalisation, via biases that do not depend on experience, reinforcement or supervision. Biases that facilitate generalisation are tuned to distinctive features that are unusual in the natural environment, such as the red and blue colours. Predispositions facilitate or hinder learning in the inexperienced brain, determining how experience is used to update the likelihood of predictive models. Neonate animals use spontaneous biases to solve the problem of induction.

## 1. Introduction

Animals must go beyond what they have experienced to respond adaptively to new challenge_1_. The ability to use sparse previous evidence in new contexts – inductive generalisation – is central to intelligent behaviour, for example in recognising an individual from a novel view-point, or predicting the behaviour or appearance of other animals. A central question is whether naïve animals, such as neonate chicks or human infants, have expectations (predispositions) for perceptual attributes including colour that can drive generalisation, or whether they build expectations by sampling their environment. To investigate spontaneous generalisation at the onset of life, we investigated poultry chicks, whose visual experience can be fully controlled from hatching until the moment of test^2, 3^. We compared observed generalisation with the predictions of an unbiased model derived from the universal law of generalisation ^4^. According to this theory, the probability that a response to one stimulus is generalized to another one depends on the perceptual distance between the two stimuli ^5^. This theory applies broadly ^6^ to different stimuli (e.g. colours, tones, shapes), sensory modalities, species and taxa (from mammals to birds, fish, amphibians and insects ^7^).

Inexperienced chicks use a fast learning mechanism called filial imprinting ^8–11^ to recognise and approach social partners after having seen them for just a few minutes, helping hatchlings to stay with the flock. Imprinting enables recognition of the mother hen, or any imprinted object, from novel points of view without explicit reinforcement ^12^. Via imprinting, chicks benefit from protection ^13–15^ and parental care ^16, 17^. Human infants are also capable of rapid generalisation from sparse evidence. For instance, 7-month infants can generalise to novel “XXY” vs “XYX” acoustic patterns, where Xs indicate identical syllables ^18^. A bit later, toddlers shown a picture of a “rhinoceros” can recognise a different rhinoceros on the television or at the zoo ^19^. Similarly, in their first hours after hatching, chicks ^20^ and mallard ducklings ^21^ use imprinting to learn the underlying structure of XX (same) vs XY (different) patterns, and generalise their affiliative responses to entirely novel XX or XY colours and sounds. Precocial birds are therefore ideal models to study spontaneous generalisation. Here we focus on colour generalisation in newborn poultry chicks (*Gallus gallus*).

Young chicks discriminate colour well ^22–24^. Chickens use colour in feeding ^25^ and social interactions ^26, 27^, including imprinting ^8, 9^. In imprinting, colour is crucial in determining approach responses ^3, 28–30^. Colour generalisation in imprinting can be controlled using visual displays ^3, 12^. In this paper, we report three sets of experiments on visual imprinting, generalisation and spontaneous preferences for colours: Experiments 1-2 compare observed generalisation behaviour to the predictions of an unbiased model after a short imprinting exposure (1-day); Experiment 3 identifies the spontaneous colour preferences present at hatching that might underlie the biases revealed in Experiments 1-2; Experiment 4 then examines the time course of learning and generalisation, looking at the effects of the biases on longer term learning (5-day imprinting exposure). We conclude by modelling spontaneous generalisation as a Bayesian process, showing how spontaneous biases (predispositions) drive learning and generalisation from sparse evidence in the neonate brain.

## 2. Spontaneous colour generalisation

We investigated spontaneous generalisation using an early exposure and generalisation test. Visually naïve chicks were first exposed (imprinted) on visual stimuli along the red-yellow (Fig. 1A) or blue-green (Fig. 1B) continua, and then tested for their approach response to the familiar stimulus vs unfamiliar stimuli from the same colour continuum.

**Fig. 1A.**
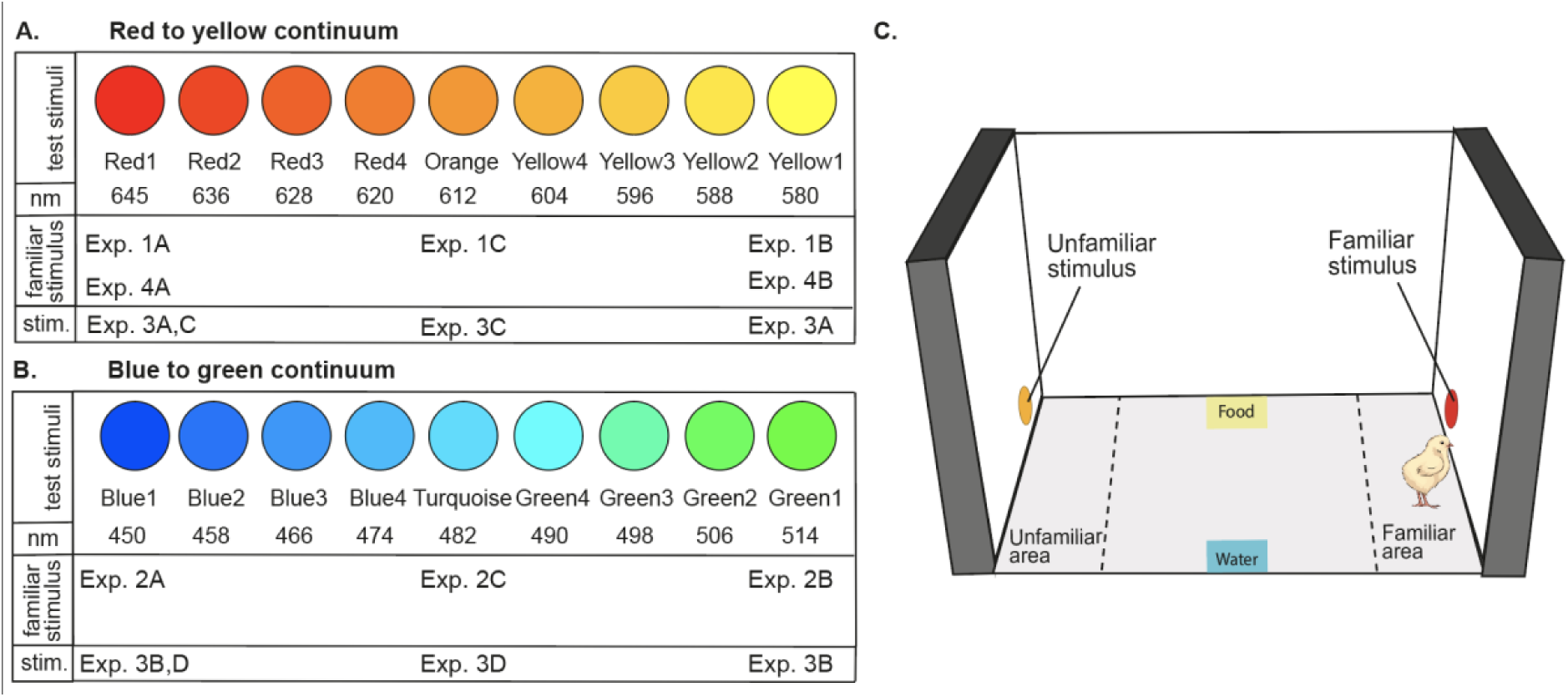
Red to yellow continuum stimuli. Colour, stimulus name and approximate wavelength (to human eye) are reported for each stimulus and experiment. Below, the experiments are listed that use a particular colour for imprinting stimulus (‘familiar stimulus’, in Exp. 1, 4) or as a choice in predisposition experiments (“stim.” for stimuli in Exp. 3AC). **B. Blue to green continuum stimuli.** Colour, stimulus name and approximate wavelength are reported for each stimulus and experiment. Below, the experiments are listed that use a particular colour for imprinting stimulus (“familiar stimulus”, in Exp. 1, 4) or as a choice in predisposition experiments (“stim” for stimuli in Exp. 3BD). **C. Controlled-rearing apparatus**. Dashed lines divide the arena into three regions: familiar stimulus area, centre, unfamiliar stimulus area.

After hatching in darkness, each chick was housed in an individual home cage (Fig. 1C) and exposed to a coloured imprinting stimulus for one day (a Ø 5 cm circle presented for 16 hours): either a red (Exp. 1A), yellow (Exp. 1B) or orange (Exp. 1C) circle on the red-yellow continuum; either a blue (Exp. 2A), green (Exp. 2B) or turquoise (Exp. 2C) circle on the blue to green continuum. The imprinting stimulus moved horizontally, switching between the right and left monitor, while the chick moved freely in the arena. A generalisation test followed with two different stimuli simultaneously presented; chicks remained in their cage and were offered the choice between a series of pairs of stimuli presented on opposite monitors (12 hours), while we measured the preference to approach the familiar stimulus (Fig. 1A and B) (see Materials and Methods for details). Trials where only the familiar stimulus was presented were used as a baseline of the strength of interest for the imprinting stimulus. The preference for the familiar stimulus was measured as:

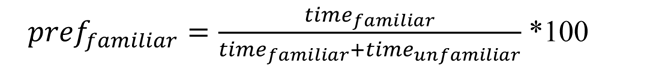

where 100 indicates a full preference for the familiar stimulus, 0 a full preference for the unfamiliar stimulus, and 50% no preference (see also ^3, 30^).

For each imprinting experiment, we compared the experimentally observed generalisation curve with the predictions of an unbiased model ^7^, derived from Shepard’s universal law of generalisation ^4^, as shown in Fig. 2. During imprinting, chicks learn the features of the imprinted object and become attached to it; they then compare it to new objects encountered, to modulate affiliative approach responses based on the perceived similarity. In an unbiased scenario, only experience is used to evaluate the similarity between familiar and novel stimuli.

**Fig. 2.**
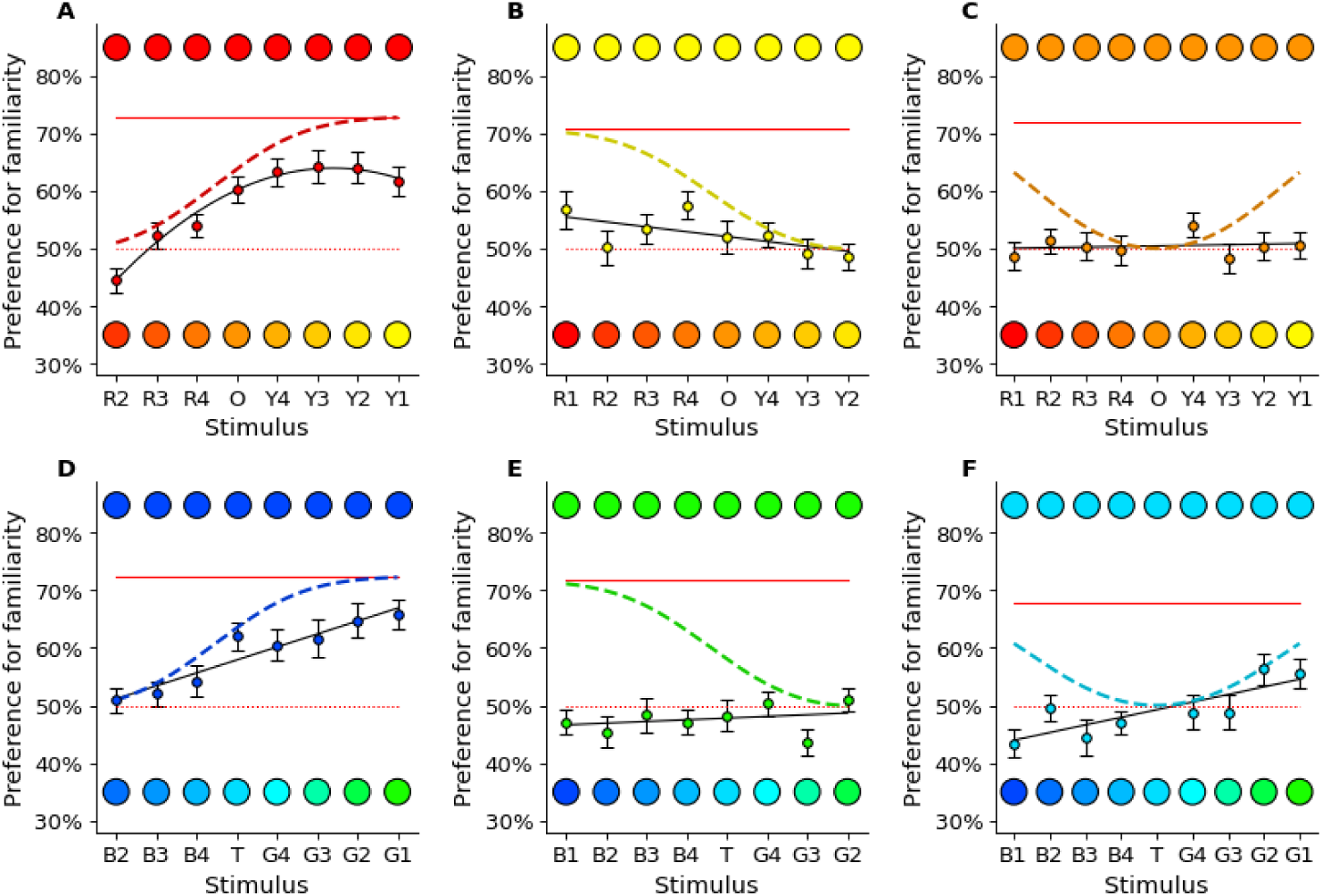
Expected and observed generalisation gradients. For each imprinting stimulus (**A**=Red1, **B**=Yellow1, **C**=Orange, **D**=Blue1, **E=**Green1, **F**=Turquoise), the predicted model for unbiased generalisation after one day of imprinting is shown with a dashed line, the observed generalisation best fit model as a solid line, with observed data as coloured dots +/-SEM. The preference for the familiar stimulus presented alone (rehearsal trials) is a red solid line, while the dotted red line indicates the absence of preference level (50%).

The unbiased model predicts a monotonic generalisation curve (with same order of stimuli in the physiological and perceptual space for humans and birds), where the preference for the familiar stimulus decreases with the distance from this known stimulus ^4, 5, 31^ (irrespective to whether we use an exponential ^4^ or a gaussian curve ^6, 7^). Although the physical and perceptual distance between stimuli along our one-dimensional colour spectrum might be different, their order is maintained in both dimensions, and the distance between two objects along the colour continuum is the same in both directions (the distance between Red1 and Red2 is the same as between Red2 and Red1). Unbiased animals are expected to display a progressive decrease of preference for more distant colours. Overall, the unbiased model predicts a consistent preference for the imprinting stimulus (no preference for the unfamiliar stimuli), a peak of preference for the imprinting stimulus, and a progressively decreasing generalisation around both sides of the imprinting stimulus (Fig. 2).

Some of the observed spontaneous generalisation curves violated the predictions of the unbiased generalisation model (Fig. 2). In Exp. 1, chicks imprinted on Red1 showed a pattern of generalisation that followed a quadratic curve (best fit), with increasing preference for familiarity as the distance between the familiar and unfamiliar stimuli increased, and a decrease in preference when the familiar stimulus was matched with the most distant colour (Yellow 1) (see Tables S1, S2). In comparing the preference to chance level, chicks had a significant preference for Red2 (t_45_=-2.59, p=0.013), and for the familiar Red1 stimulus compared to stimuli along the continuum from Red4 to Yellow1 (Red4: t_45_=2.08, p=0.044; O to Y1: t_45_>4.53, p<0.001). Conversely, chicks imprinted on Yellow1 had significant preferences for the familiar stimuli in only two cases, with an almost flat generalisation curve (see Tables S3, S4; Fig. 3E; Red4: t_42_=2.99, p=0.005; Red1: t_42_=2.04, p=0.048).

**Fig. 3.**
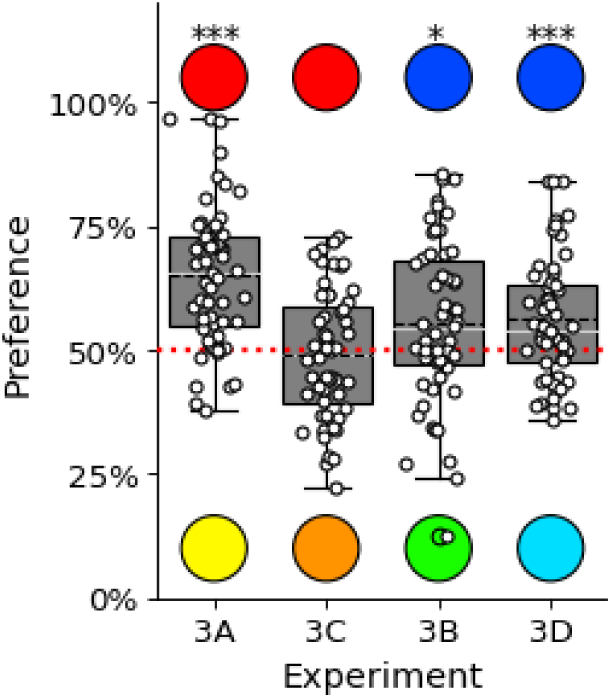
Average preference indices in the experiments 3A-3D, calculated for the predisposed Red1 and Blue1 stimuli. Boxplots display mean (black dashed line), median (white line), quartiles and outliers. Red dashed lines represent no preference (chance level); p<0.05; ** p<0.01; *** p<0.001.

This pattern was confirmed in Exp. 2. Chicks imprinted on Blue1 generalised with a gradient best fitted by a linear model (see Tables S5 and S6), indicating a linear increase in the preference for familiarity as the test stimulus increased in colour distance, with significant preferences for colours at the midpoint (Turquoise) and on the green side of the continuum (Turquoise - Green1: t_45_>3.64, p<0.001). However, similarly to what happened for Yellow1 imprinted chicks, chicks imprinted on Green1 had no generalisation gradient (see Tables S7 and S8) and only one significant preference (Green3: t_47_=-2.81, p=0.007).

In contrast to the prediction of the unbiased model, in Exp. 1A chicks preferred the unfamiliar stimulus Red2 over the familiar Red1 stimulus (Fig. 2A, t_45_=-2.59, p=0.013). This preference suggests that the chicks a predisposition to imprint on a colour for a colour less extreme on the red to yellow continuum.

In Exp. 1C and 2C, we imprinted chicks on intermediate colours Orange and Turquoise respectively. Chicks imprinted on Turquoise showed significant preferences in three cases (Green1: t_44=_2.17, p=0.035, Green2: t_44=_2.42, p=0.020, Blue4: t_44=_-2.74, p=0.009) and exhibited a linear generalisation gradient (see Table S9-S12) resembling that observed after imprinting with the colour Blue1, but with a lower intercept. Hence, the unbiased model prediction of a central peak and progressive reduction of preference with more distant colours was rejected (Fig. 2F). Similarly, chicks imprinted on Orange showed a linear flat gradient (Fig. 2C) with no significant preferences, unlike that predicted by the unbiased model.

Importantly, all chicks received the same amount of imprinting exposure in Exp. 1 ABC and Exp. 2 ABC, they were equally engaged during the imprinting phase (no significant difference in preference between any of the colours used, *F*_5, 295_=0.811, p=0.543, and had similar baseline preferences for the imprinting stimulus presented alone with a blank alternative monitor (Figure 1; Red1=73%, Yellow1=71%, Blue1=72%, Green1=72%, Orange=72%, Turquoise=68%). Therefore, the differences in generalisation between Red1 vs Yellow1 and Blue1 vs Green1 imprinted chicks are not due to different salience of the stimuli (red solid lines in Fig. 2). This shows that from the beginning of life, in the absence of previous experience, equal exposure to two stimuli along a colour continuum can have dramatically different outcomes in generalisation.

Overall, the results reject the unbiased generalisation model, indicating that spontaneous biases not dependent on reinforcement or previous experience, shape generalisation in the neonate brain. Subsequent experiments assessed whether biases in learning and generalisation correspond to early spontaneous preferences to approach a particular colour (Exp. 3, section 3), and the consequences of predispositions for the time course of learning and generalisation (Exp. 4, sections 4 and 5).

## 3. Colour predispositions and imprinting generalisation

Soon after birth, infants and domestic chicks show spontaneous preferences (predispositions^32^, see Box 1) for certain stimuli, including particular colours (see Supplementary Table 2), moving objects ^33–35^, stuffed hens ^36, 37^, hollow objects ^30^, face-like stimuli ^38^, or combinations of features such as colour and biological motion ^39^. Lesion experiments have shown a dissociation between brain areas involved in predispositions and learning^40, 41^. Whether and how predispositions drive learning and generalisation at the beginning of life is unclear. While previous work (see ^42, 43^ and references therein) has shown that colourful aposematic signals facilitate predators in learning what prey is unpalatable, the principles of spontaneous (not reinforced, unsupervised) and early generalisation are unknown. We tested whether predispositions affect unsupervised learning and generalisation in inexperienced animals.

### Box. 1. Predispositions

Predispositions are biases that influence the behaviour of animals, direct attention and approach/avoidance responses toward particular objects ^38-40^. It has been suggested that neonate vertebrates are predisposed for animacy cues, namely features associated with the presence of living beings ^38-40,45^. Predispositions orient the attention of inexperienced young animals towards specific stimuli, but it remains an open question whether predispositions or constraints influence learning and generalisation ^29^. In chicks, several predispositions to preferentially orient toward and approach some stimuli have been identified ^38,39^. Initial preferences for moving objects have been long known, and in recent years controlled studies have identified specific features of moving objects that elicit approach responses: changes in speed ^41,42^, self-propulsion ^43^, movement against gravity ^44^, biological motion ^46^. Based on the central role of moving stimuli, we imprinted chicks on moving objects. Predispositions are present also for static features, such as particular colours ^4,47-49^, for stuffed hens ^37,50^, hollow objects ^24^, face-like stimuli ^51^, and for a combination of features such as colour and biological motion ^52^. Human babies ^44-46^ and even tortoises ^47^ have similar predispositions, suggesting that they are evolutionarily conserved. Interestingly, the neural substrates of predispositions are at least partially distinct from those of learning mechanisms such as imprinting ^40^, implying that there is aseparation between the initial orienting response to “salient” objects and imprinting responses. Here, we investigate the connection between predispositions and generalisation of colour in filial imprinting.

Experiment 3 assessed colour predispositions for the colours tested in Exps. 1 and 2. If colour predispositions drive generalisation, inexperienced chicks should prefer Red1 over Yellow1, and Blue1 over Green1, that is colours that elicited fast learning and a clear generalisation gradient after imprinting. We therefore recorded colour preferences in a double choice test in the first two hours of visual exposure ^48^, at the start of the imprinting process.

Exp. 3A tested Red1 vs Yellow1, and Exp. 3B Blue1 vs Green1. As predicted, chicks preferred Red1 over Yellow1 (mean ± SEM = 0.649 ±0.018, t_58_=8.12, p<0.001, Fig. 3A) and Blue1 over Green1 (mean ± SEM = 0.554 ± 0.022, t_53_=2.40, p<0.020, Fig. 3). These preferences match the robust gradients of generalisation for chicks imprinted on Red1 and Blue1 that were observed in Exp. 1 and 2, confirming a potential role of early predispositions in generalisation.

We also assessed the preference for the intermediate stimulus vs the preferred stimulus (Orange vs Red1: Exp. 3C, and Turquoise vs Blue1: Exp. 3D), aiming to understand the shape of the initial preference curve. In Exp. 3C, chicks showed no preference for Red1 vs Orange (mean ± SEM = 0.490 ± 0.017, *t*_55_=-0.58, *p*=0.565, and in Exp. 3D chicks preferred Blue1 over Turquoise (mean ± SEM = 0.563 ± 0.017, *t*_53=_3.66, *p*<0.001, Fig. 3D). The preference for Blue1 over Turquoise suggests that the linear generalisation observed in Exp. 2C reflects a predisposition for Blue1, while the lack of preference between Red1 and Orange might explain why in Exp. 1C imprinting exposure on the intermediate stimulus (Orange) didn’t produce a generalisation gradient.

Overall, the correspondence between fast learning and generalisation after imprinting with some colours (Red1 and Blue1) but not others (Yellow1, Green1 and Orange), with early spontaneous preferences for the same colours, and the bias in generalisation (after imprinting with Turquoise), show that neonate animals like chicks are evolutionary prepared for generalisation, via biases that do not depend on reinforcement and supervision.

## 4. Time course of generalisation

To investigate how predispositions affect the time courses of imprinting and generalisation, we focused on the red to yellow continuum. Based on previous evidence (Exps. 1-3), we hypothesized that differences in generalisation curves are attributable to the chicks’ early colour predispositions. Accordingly, we used a longer (5-day) imprinting experiment on either Red1 or Yellow1 (Exp. 4A and B) to test the prediction of a theoretical model that explicitly incorporates predispositions as initial preferences and treats the imprinting process as an update of these preferences.

The model assumes that each chick has an underlying preference for each stimulus and chooses between them in proportion to their evaluated value:

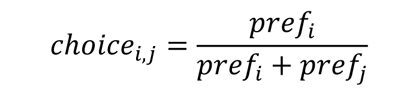

where *f*_*i*_ and *f*_*i*_ represent the value the chick assigns to the stimulus, and *choise*_*ij*_ indicates the proportion of choices of stimulus *i* over stimulus *j*. When comparing the model predictions to empirical observations, we scaled the model predictions to the maximum observed preference (either in rehearsal trials or during the tests).

During imprinting, chicks use their experience to re-evaluate and adjust their preferences. The learning process is represented here as a gaussian and multiplicative update of the preferences. At each time step *t*, preferences across the stimulus continuum are updated as:

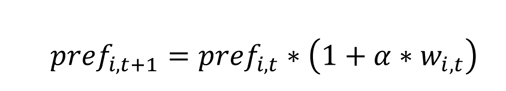

where the update rate *α* is a constant that defines the speed of learning, and the update weight function *w* describes the experience of the chick. It is maximal at the stimulus value presented at time *t*, and has a narrow gaussian distribution around the peak. Preferences for stimuli that are perceptually very similar to the exemplar are updated as well. At each time step, the preference curve is normalised to its maximum.

We applied this model to the red-yellow continuum (Fig. 4), using the initial preference curves estimated from Exps. 3A,C, where Red1 and Orange were equally preferred at hatching, and approximately twice as attractive as Yellow1. In choosing Red1 and Yellow1 as the imprinting stimuli, we iterated the update process for 400 time steps, setting *α* = 0.01 and the update curve’s standard deviation to *σ* = 0.2, and calculated the preference curves at each time step. The model predicts that preferences for the predisposed Red1 and the non-predisposed Yellow1 develop differently (Fig. 4). By the end of the simulation the imprinting stimulus is strongly preferred for in both cases, but the preference curves differ at intermediate time steps. For the predisposed Red1, the preference forms quickly, and the imprinting narrows it over time. In contrast, for the non-predisposed Yellow1, establishing the preference takes a longer (imprinting) time, with an intermediate phase when all stimuli on the red-yellow continuum are expected to be equally attractive.

**Fig. 4.**
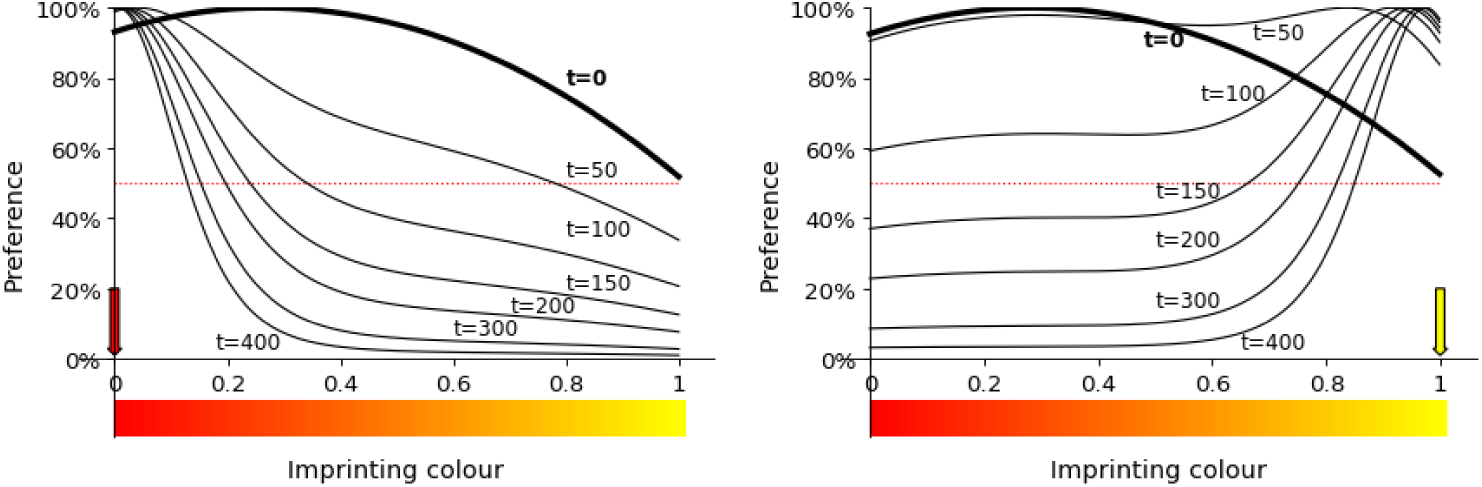
Predicted colour preferences over time. The preferences develop differently when imprinting on Red1 **(A)** and Yellow1 **(B)**, despite converging onto the same strong preference for the imprinting stimulus. The red and yellow arrows highlight the imprinting stimulus, and ‘t’ indicates time step.

Experiment 4 tested the model empirically on the red-yellow continuum, comparing its predictions to the results of the 1-day imprinting experiments (Exp. 1A, 1B) and running a new, longer experiment, where generalisation is tested after five days of imprinting, when the learning is completed and stable^49^. We used the same imprinting procedure, generalisation test and analyses as for the 1-day imprinting experiments (Exp. 1 and 2), the only difference being that the imprinting exposure was repeated for five consecutive days before the generalisation test.

We found that the model reflects the results of the short 1-day imprinting (Exp. 1A, 1B) and the long 5-day imprinting experiments (Exp. 4A, 4B) (Fig. 5). For the short (one day) imprinting experiments, the model predicts a gradually increasing preference for the familiar stimulus when imprinted on Red1 (Fig. 5A) and a slight preference for Yellow1 over unfamiliar stimuli. After the long (5 days) imprinting (Exp. 4), the predicted generalisation curves have similar shapes for both imprinting colours: generalisation to the most similar colour, and then a sharp increase of preference to a plateau. The model captures most of the variation in the experimentally observed generalisation curves: Exp. 1A, short imprinting on Red1: R^2^=0.979; Exp. 4A, long imprinting on Red1: R^2^=0.807; Exp. 4B, long imprinting on Yellow1 R^2^=0.738. For Exp. 1B, short imprinting on Yellow 1, the trend in the observations is small and accordingly the R^2^ value is low, 0.392. We conclude that accounting for the initial preferences, and treating the imprinting as an update of these initial preferences, is sufficient to explain the different generalisation curves and time courses for different colours. This conclusion underlines the importance of predispositions in learning from the first day of life, in the absence of any reinforcement and previous experience.

**Fig. 5.**
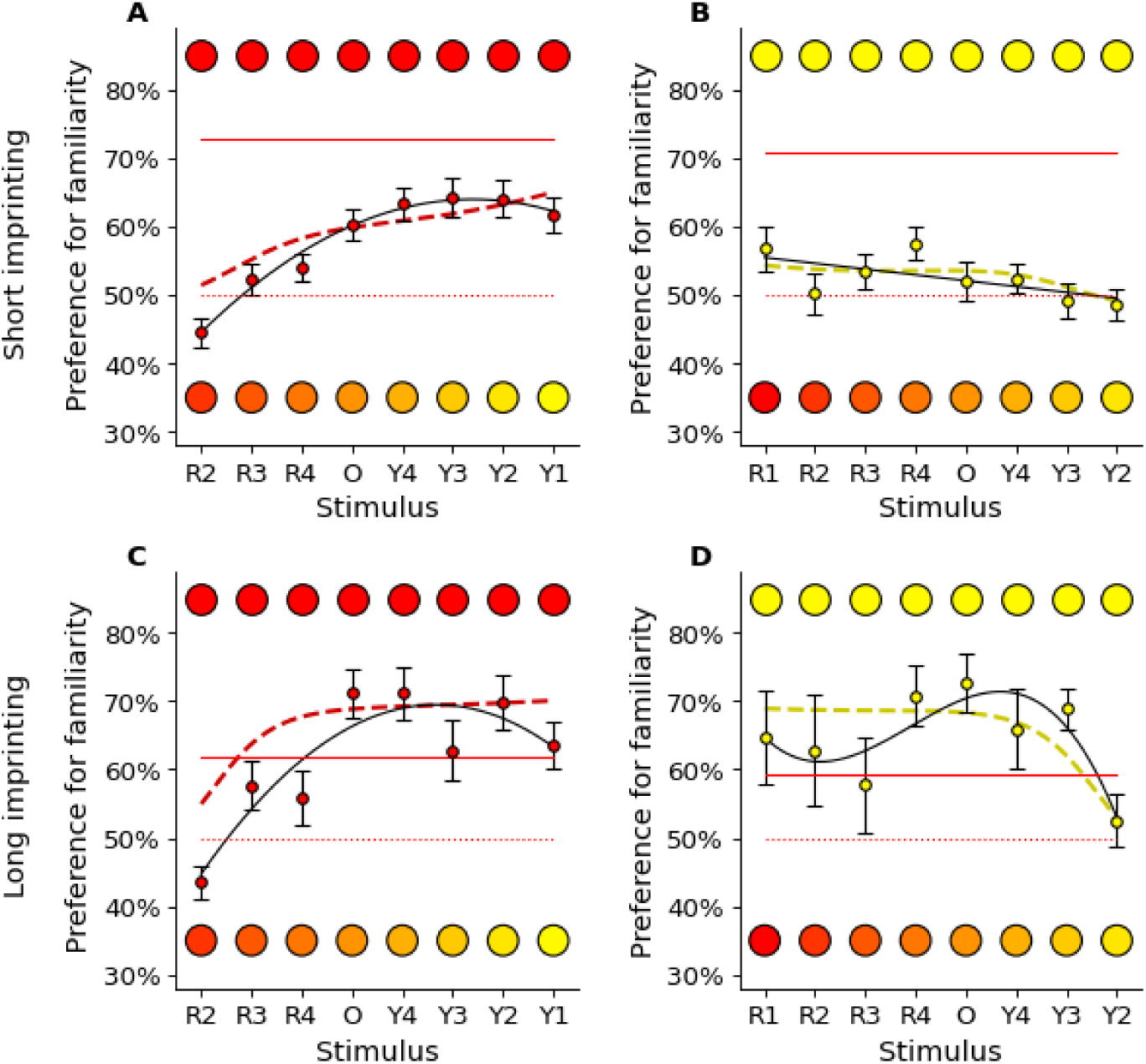
Experimentally observed vs predicted generalisation curves after short (1-day; t=100) and long (5-day; t=300) imprinting experience, for Red1 (**A**, **C**) and Yellow1 (**B**, **D**) imprinting stimuli (Exps. 1, 4). For each imprinting stimulus and duration of imprinting, model predictions are shown with a dashed red line, the observed generalisation best fit model as a solid line, and the observed data as coloured dots +/-SEM. The preference for the familiar stimulus presented alone (rehearsal trials during Exps. 1,4) is a red solid line, while the dotted red line indicates the absence of preference. The predictions of the theoretical model match the empirical observations. Note that in the long experiments (Exp. 4), the rehearsal trial preference is lower than preference in the dual choice experiments (after the end of the sensitive period, chicks actively avoid the unfamiliar stimulus when an option is given).

## 5. Bayesian conditional probabilities drive the evolution of predispositions

It is not clear how predisposed preferences help chicks to identify the correct imprinting target in a generalisation task, when alternatives are available. Several studies have focused on the similarity between predispositions for specific features (e.g., specific colours or movement dynamics^35, 39, 50–52)^ and the characteristics of the mother hen. However, preferences for different colours vary between experimental settings (reviewed in Supplementary Table 2), suggesting that the physical characteristics of the stimuli (e.g., a specific wavelength) are not sufficient to capture observed behaviour. We use a different approach, focusing on the relative preferences, considering the task similar to a likelihood test. We formalise this idea modelling the chick as an ideal Bayesian observer that evaluates each potential target according to the conditional probabilities calculated with Bayes’ theorem:

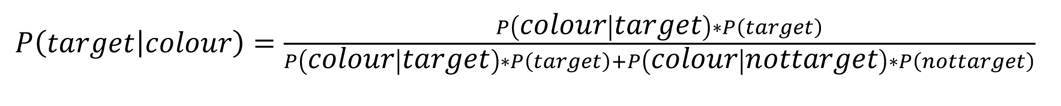

where *P*(target|*colour*) is the probability that the chick assigns to the hypotheses that an object seen is the target (the mother hen), given its observed colour. In our formulation we assume that the chick hatches with a set of expectations (predispositions), both about the probability distribution of the features of the ideal target *P*(*colour*|target), and about the probability distribution of the features of other objects *P*(*colour*|*not-t*-target). We will treat the probability that any given object is the target *P*(target) or something else *P*(*not-target*) as constants. It follows that the likelihood ratio of an object with a given colour being the target, and therefore the interest the chick should show in it, will increase with the probability that the target displays that particular colour *P*(*colour*|*target*), and decrease with the prevalence of the colour in other objects present in that environment *P*(*colour*|*not-target*).

From an evolutionary point of view the ideal imprinting targets for non-domesticated chicks, the mother hens, display yellow and brown colours with reddish areas in the head region (Fig. 6B ^53^). In a previous study, we investigated the feather and head colours of red junglefowl hens (the specimens closest to ancestral chickens before domestication), using spectral reflectance measurements of bird skins ^53^. The reflectance curves of the feathers show a gradual increase with larger wavelengths (characteristic of phaeomelanin), giving the specimens their yellowish-brown appearance. We have now calculated the avian photoreceptors’ responses to these curves and plotted the results in the tetrahedral colour space of birds (Fig. 6C). The feather colours of various areas of red junglefowl hens fall close the achromatic middle point, except for the lack of UV. In addition to the feathers, the soft parts of the head regions (comb, eye) appear reddish ^53^. Hence, red junglefowl hens are equally likely to display reddish and yellowish hues, and the coloration of the mother hens does not explain the chicks’ preference for red in itself.

**Fig. 6.**
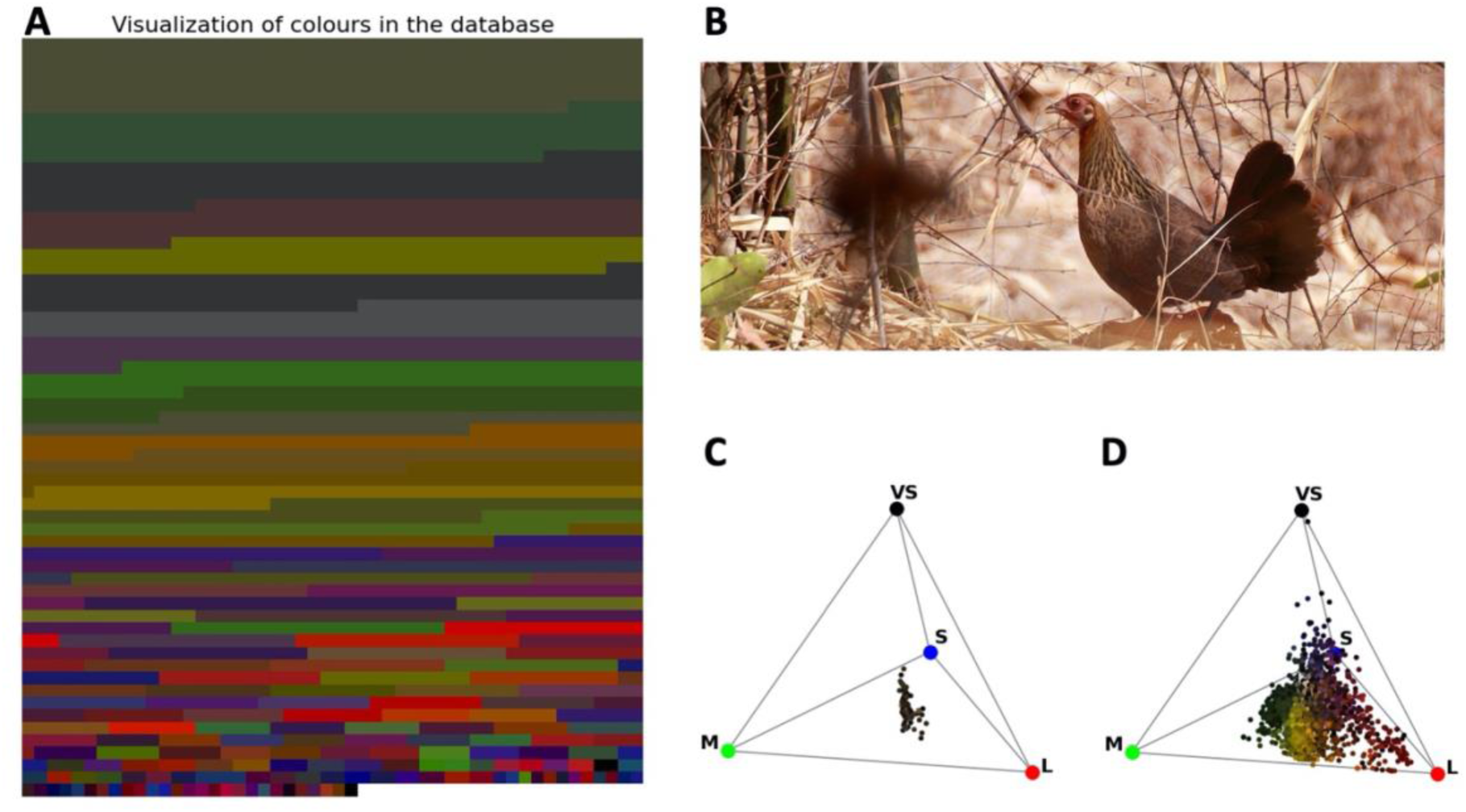
(A) The “colours” of the world. We calculated the red, green and blue photoreceptor responses of a typical UV-sensitive bird _56_ to a large library of natural colours ^54, 55^ and visualised them as RGB values. The area of each coloured stripe corresponds to its frequency in the database. Natural objects – flowers, leaves, rocks, soil – appear as unsaturated greens, browns and greys to the birds. Red coloured objects exist, but they are rare. Note that the colours have been normalised for equal total brightness, and that the visualisation ignores the UV dimension of colour. **(B) The red junglefowl hen** (the closest relative of the domestic chicken) is predominantly yellow/brown, with reddish areas in the head region. Photo credit: Brian Gratwicke, downloaded from https://commons.wikimedia.org/wiki/File:Red_Junglefowl_hen_India.jpg. **(C) The colour loci of the red junglefowl** hen feathers ^53^and of **(D) natural objects**, plotted in the tetrahedral colour space of birds. The tips of the tetrahedron correspond to situations where only one receptor is activated. VS – UV/violet sensitive receptor, S – short wavelength (blue) sensitive receptor, M – medium wavelength (green) sensitive receptor, L – long wavelength (red) sensitive receptor.

Turning to the colours of the natural environment, we used spectral reflectance functions of a variety of materials including soil, minerals, vegetation and flowers, available in public databases (FReD^54^ and USGS^55^; see Methods). Plotting the avian photoreceptor responses ^56, 57^ in tetrahedral space ^56^, reveals a lack of saturated colours, and that red is rare while greens/browns/dark yellows dominate (Fig. 6). Against such a background, red junglefowl plumage colours would be hard to identify, but the reds of the soft parts on the head would stand out. Therefore, as there is a larger probability that a hen (target) displays a red colour *P*(*colour*|*target*) (even though it is primarily brown), and a smaller probability that a non-target object in the environment is red *P*(*colour*|*not-target*), we conclude that the colour red is a good indicator for identifying the target mother hen *P*(*target*|*colour*). For this reason, red (and not yellow) colour can be used by newly hatched chicks initially for orienting towards the mother hen, and then recognise the target compared to alternative objects, making red colour an optimal target for early predispositions. Therefore, Bayesian conditional probabilities offer an explanation for the red over yellow preference we have observed in Exp. 1, 3 and 4.

## 6. General discussion: rara avis in terris

To cope with environmental changes and novelty, it is necessary to go beyond previously observed data and solve the *problem of induction*, making connections between previous observations and expectations or predictions about what has not yet been observed ^58^. In recent decades it has emerged that brains work by making predictions, rather than simply by reacting to sensory stimuli ^59^. After having seen only white swans, one might predict that the next swan will be white. This reasoning does not apply only to birds. The Roman poet Juvenal, almost two thousand years ago, referred to a *rara avis in terris, nigroque simillima cygno*, “a bird as rare as a black swan”, to indicate an exceptional person, with features that are not expected or predicted. But where do expectations come from? Generalisation is especially demanding when experience is limited, as at the beginning of life or in any new context. Infants ^18^, children ^19^ and young animals ^20^ generalise effectively from sparse evidence, but the strategies and mechanisms are unknown. This knowledge gap derives from the difficulties of testing naïve individuals whose experience is controlled from birth to test. Ethical and practical issues prevent and limit controlled-rearing studies in human babies and altricial species. To address this issue, we investigated the principles of spontaneous generalisation in neonate and visually inexperienced chicks, which are a precocial animals hatched with a mature sensory and motor system. Chicks were controlled-reared from hatching to test in an arena with coloured displays. We investigated learned and spontaneous approach responses to coloured moving objects, which young chicks treat as social partners, and compared empirically observed generalisation (choice between a familiar and several unfamiliar shapes with colours progressively more distant from the familiar ones) with the predictions of an unbiased generalisation model ^61^.

The results reject the unbiased model ^4^, implying that the predispositions ^32^ that orient first approach responses facilitate inductive generalisation in neonate chicks. First, inconsistent with unbiased predictions, chicks exposed and then tested for generalisation on the red to yellow and blue to green continua exhibited completely different generalisation curves, depending on their imprinting experience (Red1, Yellow1, Orange or Blue1, Green1, Turquoise in Exp. 1ABC, Exp. 2ABC respectively). Chicks had been exposed to each imprinting stimulus for the same amount of time, and were equally responsive to each stimulus during imprinting. Despite this, only those imprinted on Red1 and Blue1 exhibited a robust generalisation gradient, while chicks imprinted on Yellow1, Orange and Green1 showed no preference for either the familiar or unfamiliar stimuli, and chicks imprinted on Turquoise did not show the expected peak preference for the familiar stimulus. These differences in generalisation after similar experience are incompatible with an unbiased model, and point towards an evolutionarily-prepared model of generalisation, in place at birth.

We asked whether learning and generalisation to novel objects are driven by early predispositions to approach different colours. Predispositions to approach different colours in the first visual experience (Exp. 3ABCD) matched the results of the generalisation test: chicks spontaneously preferred Red1 over Yellow1, Blue1 over Green1, and Blue1 over Turquoise, but had no preference for Red1 vs Orange. This corresponds to a scenario where neonate chicks operate as Bayesian observers ^31^. In fact, red and blue colours are disproportionately present in objects that are likely to be the imprinting target, rather than in background objects, as shown by our analysis of fowl and environmental colours^53–55^ (section 5).

Second, we observed preferences for the unfamiliar stimulus Red2 (Exp. 1 and 4), whereas the unbiased generalisation model ^28, 31^ predicts a preference for the imprinting object (Red1). A preference for unfamiliar stimuli^20^ or for stimuli “slightly different” ^48^ from the imprinting object has been repeatedly observed in chicks. The preference for unfamiliar stimuli was initially observed at the beginning of the imprinting process ^48^, and described as the outcome of two complementary components: the attraction for familiar objects and that for novel objects. This dual process would allow young birds to gain as much information as possible about their specific mother and siblings from the different view-points, to enhance subsequent recognition. However, this explanation cannot account for why preferences for unfamiliar objects have been observed also in older chicks, after several days of imprinting exposure ^20, 32, 60^, and the selective presence of novelty preferences that we observed only for certain novel objects. This pattern suggests that the preference for unfamiliar stimuli might be due to initial biases for specific colours, and in particular that Red1 is not the most initially preferred colour for chicks in the Red1 to Yellow1 continuum. This is supported by the observation that Red1 and Orange were equally attractive to naïve chicks (Exp. 3C). The preference for colours different from the imprinting stimulus observed after one day, and even five days of imprinting, supports the hypothesis that spontaneous, naturally-selected biases influence chicks’ learning and generalisation: they have prepared minds. These biases are likely to reflect the evolutionary significance of particular colours. A predisposition to approach red has a selective advantage, because the head region of the red jungle fowl hens, the closest relatives to ancestral chickens before domestication, is reddish^53^. Hence, a predisposed preference for red could help chicks to selectively process and learn from reddish stimuli, that are associated with the target imprinting stimulus.

Overall, we show how spontaneous biases (predispositions) for behaviourally relevant stimuli can facilitate learning with limited experience, leading to faster and more robust generalisation, compared to generalisation from less predisposed stimuli. This finding has broad implications for understanding the development of cognition in biological and artificial minds. Predispositions not only guide early orienting approach responses but they directly affect learning and generalisation, showing that the neonate brain makes predictions consistent with Bayesian models^31, 61^, in the absence of previous experience and reinforcement. Understanding these biases could contribute to the design of educational materials and interventions for infants and children with typical and atypical development.

Moreover, the connection between early predispositions and generalisation suggests that low level biases or Bayesian expectations can be implemented as building blocks of more sophisticated cognitive abilities, with the potential to improve artificial intelligence, that currently requires extensive training or preparation ^62^. AI may benefit from priors or biases similar to those used by infants, to more efficiently process and learn evolutionarily-or task-significant information. By identifying the biases that influence an animal’s learning, researchers may be able to design systems that can quickly adapt to novel environments, and to understand the building blocks of cognition.

## 7. Methods

### 7.1 Incubation and rearing conditions

Fresh eggs of domestic chicks (*Gallus gallus*) Ross 308 were used in all experiments. Eggs were incubated and hatched in darkness under controlled conditions (37.7°C and 40% humidity). Chicks hatched in opaque individual boxes, and had no visual experience before the experiment. Within 8-18 hours from hatching, chicks were sexed and individually placed in their home cage with water and food available ad libitum. Chicks were exposed to a day:night cycle of 16:8 hours, alternating stimuli (day) and dark slides (night) displayed on the screens.

### 7.2 Apparatus

The home cage was used as experimental apparatus. Apparatuses were rectangular (90 x 60 x 60 cm) enclosures, with white walls and a computer located on each of the short sides of the arena (Figure 1C). Stimuli were displayed on high-frequency 24-inch widescreen monitors (ASUS MG248, 144 Hz). A webcam located above the centre of the arena recorded chicks’ behaviour. Food and water were placed in the middle of the long walls, available ad libitum. The areas within 20 cm of each monitor were defined as the regions near familiar/unfamiliar stimulus. These areas were chosen as chicks spent above 90% of their time in these regions (average across all chicks from Exp. 1A).

### 7.3 Imprinting: Experiment 1,2 and 4

#### 7.3.1 Subjects

We tested the following number of chicks: Exp. 1A: 24 males, 24 females; Exp. 1B: 24 males, 24 females; Exp. 1C 28 males, 31 females; Exp. 2A: 24 males, 24 females; Exp. 2B: 24 males, 24 females; Exp. 2C 26 males, 24 females; Exp. 4A 15 males, 13 females; Exp. 4B: 15 males, 20 females.

#### 7.3.2 Stimuli

Stimuli were circles of different colour with a diameter of 5 cm, including a 1.5 mm black outline, presented on computer monitors. Previous experiments showed the efficacy of similar visual displays as imprinting objects^20, 32, 52, 53^. The approximate wavelengths for all test stimuli are shown in Figure 1A and B, with corresponding RGB values listed in Supplementary table 1. In each experiment we presented one imprinting stimulus and eight test stimuli. The stimuli move horizontally at a speed of 10 cm/s, with a frame rate of 120 fps. As the stimulus reached one end of the screen, it disappeared out of the frame for 0.5 seconds before reappearing. Videos were displayed using Potplayer (1.7.17508).

#### 7.3.3 Procedure

The experiment consisted of two phases: imprinting and test phase. In Exp. 1 and 2, the imprinting phase took place on day 1, in Exp. 4 on day 1-5, During the imprinting phase, chicks were exposed to a single imprinting stimulus that was displayed on one of the screens. The side of the display screen was counterbalanced. Each chick underwent 16 hours of imprinting (8 x 2-hour imprinting sessions). We focused the analyses on the first 12 hours, since in Exp. 1 we noticed that in the last 4 hours chicks spent a high proportion of time asleep. Each imprinting session consisted of 10 trials, with 10 minutes of stimulus presentation in each trial, followed by 2 minutes of empty white screen.

The test phase took place either on day 2 (Exp. 1 and 2) or on day 6 (Exp. 4) after hatching. The test phase consisted of eight sessions of 10 trials (10 minutes of stimulus presentation followed by 2 minutes with a white screen). Each session began with two rehearsal trials which were identical to the imprinting trials (only one imprinting stimulus was presented on one monitor). This was followed by eight test trials, in which chicks were exposed to two stimuli, each presented on a different monitor: the imprinting stimulus and one of the test stimuli. A different test stimulus was presented in each trial. Therefore, each chick experienced all eight test stimuli in one session.

#### 7.3.4 Data analysis

Statistical analyses were performed using R (version 4.0.4), packages ply4, lme4, ez, tidyverse, ggplot2, and gridExtra and Google Colab using Python’s scipy and statsmodels packages. Alpha level was set to p≤0.05. P values have been corrected for multiple comparisons using Bonferroni-Holm correction.

The position of chicks’ and stimuli’s centroid during the experiments were analysed using the automated tracking software DeepLabCut^63^. We analysed only frames tracked with a likelihood reliability of 0.9 or above ^52^: in Exp. 1 and 2 this corresponded to 98% of recorded frames, in Exp. 4 to 99%. The preference for the familiar stimulus was measured as:

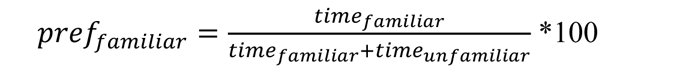

where a preference index of 50% indicated no preference for either stimulus, a preference higher than 50% indicated a preference for the imprinting stimulus, and a preference lower than 50% is preference for the novel stimulus (see also ^3, 30^).

To assess the chicks’ preference for the imprinting stimulus during the imprinting phase we ran mixed-model ANOVA with preference scores as the dependent variable, Sex (male, female) and Stimulus as the independent variables. One-sample t-tests were used to assess whether chicks spent significantly more time near the imprinting stimulus than chance level (50%). For the test phase, a two-way mixed-model ANOVA was run to assess the effect of Stimulus on chicks’ preferences for familiarity, with the percentage of time spent near the imprinting stimulus as dependent variable and Sex (male, female) and Stimulus as independent variables. Greenhouse-Geisser correction was applied to within-subjects factors that violate the sphericity assumption. We used one-sample t-tests to assess whether chicks spent significant different time near the imprinting stimulus than chance level (50%) when presented with each novel stimulus. Independent samples t-tests were used to assess whether chicks showed significantly different preferences towards the imprinting stimulus when presented with a test colour from the continuum half on either side of the intermediate imprinting stimulus.

### 7.4 Experiment 3: Colour predispositions

#### 7.4.1 Subjects

In Experiment 3, we used 256 domestic chicks: Exp. 3A: 34 males, 25 females; Exp. 3B: 25 males, 29 females; Exp. 3C 25 males, 31 females; Exp. 3D: 23 males, 31 females.

#### 7.4.2 Stimuli

We used a subset of the stimuli described in 3.2 and illustrated in Fig. 1A. In Exp. 3A we tested the preference for Red1 vs Yellow1, in Exp. 3B we tested the preference for Blue1 vs Green1, in Exp.3C Red 1 vs Orange, in Exp. 3D Blue1 vs Turquoise.

#### 7.4.3 Procedure

After hatching in darkness, chicks were sexed and individually placed in their apparatus. Each chick underwent six consecutive trials of double choice test, with one stimulus presented on each screen. Each trial consisted of 20 minutes of stimulus presentation, followed by 2 minutes of white screen. The right-left side on the stimuli was counterbalanced between trials.

#### 7.4.4 Data analysis

We used two measures of spontaneous preferences were used: the first stimulus approached and the preference scores for the Red1 or Blue1 stimulus (depending on the experiment). The first stimulus approached was defined as the first choice area entered by the centroid of the chick. Preference scores were calculated as described in Exp. 1 and 2. Binomial tests were performed to analyse the number of first approaches toward each of the test stimulus. A two-way mixed-model ANOVA was used to assess the percentage of time that the chicks spent close to each stimulus, with the preference score as the dependent variable and sex (male, female) and trial (1 to 6, within subjects) as the independent variables. One-sample t-tests were used to assess whether chicks spent significantly different time near each stimulus than chance level (50%).

### 7.5 Computational methods

#### 7.5.1 Calculating avian photoreceptor responses to colours

Avian photoreceptor responses were calculated for the feather colours of red junglefowl hens and for a library of natural spectra with the aim of relating the experimentally observed colour predispositions to the colours of the correct imprinting target (the mother hen) and the colours of the environment. Following ^64^, the photoreceptor quantum catches of an animal can be calculated using the equation:

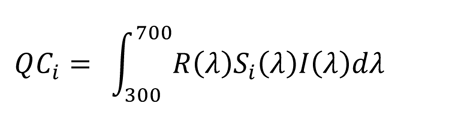

where *QC_i_* is the camera quantum catch of the sensor *i*, *R* is the spectral reflectance function of the stimulus, *S_i_* is the spectral sensitivity function of the receptor *i*, and *I* is the photon flux of the illuminant, in this case set to 1 across all wavelengths. We used the receptor sensitivity functions of an average ultraviolet-sensitive avian viewer ^56, 57^ to calculate the quantum catches in response to our measurements of the hen feather reflectances ^53^ and for the natural objects from two publicly available databases. The flower reflectance database (FReD ^54^) is a large database of flower and leaf reflectance data, consisting of 2,494 spectra. 536 additional spectral reflectance functions of various types of vegetation, soil and minerals were extracted from the spectral library of the U. S. Geological Survey (USGS^55^).

The quantum catches of the four (ultraviolet, blue, green and red) colour cones *QC_i_* were normalized to sum to 1, yielding relative {vs s m l} values. These {vs s m l} values were then projected to the tetrahedral colour space of birds, following the equations from ^56^:

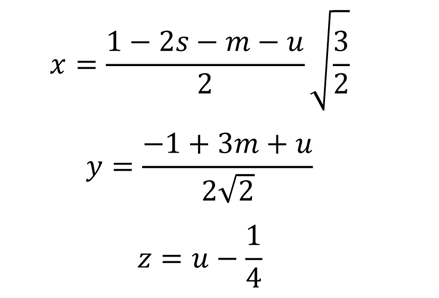

where x, y, and z are Cartesian coordinates in three-dimensional (3D) space. If a colour stimulates only one class of photoreceptor, then its point (x, y, z) lies at the appropriate vertex (tip) of the tetrahedron; when all four cones are stimulated equally, in other words, the colour is achromatic, the point is at the origin (0,0,0).

## Acknowledgements

This work was supported by the Leverhulme grant RPG-2020-287 (EV) and the Royal Society Leverhulme Trust fellowship SRF\R1\21000155 (EV).

## Authors and Affiliations

1. School of Biological and Behavioural Sciences, Queen Mary University of London, London, UK
2. School of Life Sciences, University of Sussex, Brighton, UK
3. Alan Turing Institute, London, UK

## Contributions

SW, VV, EV designed research; SW, LF, VV performed the experiments; SW, LF, VV analysed the data; SW, VV, DO, LF, EV contributed to methodology; SW,VV plotted data/results; SW, VV, EV wrote the paper with input from LF and DO; SW, VV, LF, DO, EV reviewed and edited the paper; EV provided funding and supervision.

## Competing interests

The authors declare no competing interests.

---

**Table S1.** Colour name, RGB values and related approximate wavelength (nm) of the stimuli used in our experiments.

| Colour | R | G | B | Approx. nm |
| --- | --- | --- | --- | --- |
| Yellow1 | 255 | 255 | 0 | 580 |
| Yellow2 | 255 | 230 | 0 | 588 |
| Yellow3 | 255 | 203 | 0 | 596 |
| Yellow4 | 255 | 176 | 0 | 604 |
| Orange | 255 | 148 | 0 | 612 |
| Red4 | 255 | 119 | 0 | 620 |
| Red3 | 255 | 87 | 0 | 628 |
| Red2 | 255 | 52 | 0 | 636 |
| Red1 | 255 | 0 | 0 | 645 |
| Green1 | 26 | 255 | 0 | 514 |
| Green 2 | 0 | 255 | 70 | 506 |
| Green 3 | 0 | 255 | 169 | 498 |
| Green 4 | 0 | 255 | 255 | 490 |
| Turquoise | 0 | 222 | 255 | 482 |
| Blue4 | 0 | 187 | 255 | 474 |
| Blue3 | 0 | 151 | 255 | 466 |
| Blue2 | 0 | 113 | 255 | 458 |
| Blue1 | 0 | 79 | 255 | 450 |

**Table S2.** Overview of previous spontaneous colour preference experiments in precocial birds. Adapted and expanded from Ham and Osorio (2007). Extended references are provided under the table.

| Species and strain | Age | Stimuli | Behaviour | Colour preferences | Reference |
| --- | --- | --- | --- | --- | --- |
| White Rock chicks (*Gallus gallus*) | 1 day | Coloured Ostwald chips | Pecking | Preference for orange and blue over yellow and green | Hess (1956) |
| Pekin ducklings (*Anas platyrhynchos domestica*) | 1 day | Coloured Ostwald chips | Pecking | Preference for green and yellowish-green over orange and blue | Hess (1956) |
| White Rock chicks (*Gallus gallus*) | 13-16 hours | Coloured spheres | Approach | Order of preference: orange, green, red, blue, yellow | Schaefer, Hess (1959) |
| White Rock chicks (*Gallus gallus*) | 30-36 hours | Coloured spheres | Approach | Order of preference: orange, red, green, blue, yellow | Schaefer, Hess (1959) |
| Domestic chicks (*Gallus gallus*) | 18-30 hours | Coloured balls | Approach | Equal preference for red, green, white | Smith, Hoyes (1961) |
| White Leghorn chicks (*Gallus gallus*) | 12-24 hours | Rectangle, outline of hen | Time spent | Preference for red over white | Smith, Meyer (1965) |
| White Leghorn chicks (*Gallus gallus*) | 1 day | Coloured chicks | Approach | Equal preference for red, yellow, green | Salzen, Cornell (1967) |
| White Leghorn chicks (*Gallus gallus*) | 8 days | Coloured chicks | Approach | Preference for yellow and green over red | Salzen, Cornell (1967) |
| Cobb chicks (*Gallus gallus*) | 2 days | Coloured pen | Approach | Preference for red and yellow over blue | Taylor, Sluckin, Hewitt (1969) |
| Japanese quail (*Coturnix coturnix*) | 4-7 days | Coloured pen | Approach | Equal preference for red and blue | Taylor, Sluckin, Hewitt (1969) |
| White leghorn chicks (*Gallus gallus*) | 1 day | Coloured paper disc | Approach | Preference for red over yellow or green | Salzen et al. (1971) |
| Rhode Island red chicks (*Gallus gallus*) | 1 day | Flashing stimulus patterns | Approach behaviour | Initial aversion to green; equal preference for white, blue, yellow, and red | Kovach (1971) |
| White leghorn chicks (*Gallus gallus*) | 1 day | Coloured light | Pecking | Preference for orange-red and blue-violet | Fischer et al. (1975) |
| Bobwhite quail chicks (*Gallus gallus*) | 1 day | Coloured food | Pecking | Preference in order: blue, green, yellow, red | Mastrota, Mench (1995) |
| ROSS 308 chicks (*Gallus gallus*) | 1 day | Moving circles on computer monitors | Approach preferences (time) | Preference for red over yellow | Freeland et al. (2023) |

## Notes

### Competing Interest Statement

The authors have declared no competing interest.

